# The Updated Properties Model (UPM): A topology optimization algorithm for the creation of macro-micro optimized structures with variable stiffness

**DOI:** 10.1101/2022.10.21.513158

**Authors:** Luis Saucedo-Mora, Ismael Ben-Yelun, Hugo García-Modet, Miguel Ángel Sanz-Gómez, Francisco Javier Montáns

## Abstract

The design and manufacturing of high value industrial components is suffering a change of paradigm with 3D printing. In this change of paradigm, metamaterials have an important role because when a component is 3D-printed, it is performed from the micro level, where custom structures may be designed to endow the material and the component of special of customized mechanical properties. Topology optimization techniques facilitate the design of both the microstructures and the overall component topology, and today the component topology may be designed assuming a continuous spectrum of mechanical properties facilitated by different locally designed microstructures. However, current topology optimization techniques do not operate directly with the mechanical properties of the material, but through density intermediates, using density-based limits like a minimum or maximum density, assuming an homogeneous base material. We propose here a novel topology optimization algorithm which operates directly on the mechanical properties and energies, without employing density intermediates. The proposed approach reduces the algorithmic complexity since the optimization is performed by the direct iterative update of the mechanical properties, through information taken from its finite element analysis. We show that the proposed methodology can reach similar results as the current techniques based on a gradient descent optimization, eliminating the need for external parameters and, hence, increasing the easy of use and its robustness. The proposed technique is specially suitable for two-level concurrent material-component design using functionally graded metamaterials.

## 1. Introduction

When a material is part of a structure, there is a need to consider the whole as an indivisible entity, because its integrity depends on a good structural design and a proper behaviour of the material along time, bearing in mind that all the processes of degradation and damage of the structure during its service life arise in the microstructure of the material.

As new challenges, the society demands lighter and more durable structures with materials that are increasingly more advanced. A join optimization process of the material and the structure often pursues designs inspired by nature, leading to a more efficient use of resources. When a biological element grows to form a structure, such as a bone or a tree, it is done through a process called morphogenesis [1], in which the material and the structure are combined and adapted simultaneously to achieve a functional and durable element with the minimum resources and optimal properties for the given external actions. One of these mechanisms is the ontogenetic adaptation [2] where the deformation of the material is the key agent to define the growth of the structure. It is capable to adapt an optimal design to the external variable conditions through a combination of changes in both; the shape of the structure and the microstructure of the material. An example of the close relationship between the material and the structure in natural optimal designs is explained by the Wolff’s law [3]. It defines the adaptation of the bones when there is a change of its external loading conditions, as in the case of prosthesis in a joint. Many biological structures are made of functionally graded microstructures that endow the continuum with different mechanical properties using frequently the same base material.

The search for optimal component designs made of optimal materials has been a constant in human evolution. During industrial evolution, the development of optimal goods has been driven by the parallel development of three factors: the development of materials, the development of manufacturing procedures, and the development of techniques to calculate the optimal design. The availability of 3D printing is a change of paradigm in the design of industrial components because it eliminates many of the manufacturing constraints and, given that components are printed from the microscale and mesoscales, it brings the possibility of designing the material itself through the design of the microscale. These designs constitute a material in the classical sense when seen at the continuum scale, and are called metamaterials. These metamaterials have designed or unique mechanical (or other) properties. With 3D printing, a component may have a spectrum of mechanical properties from designed functionally graded metamaterials. In contrast to classical materials, the same printing base material results in radically different continuum properties through a suitable microscale printed design. Mechanical properties like the Young modulus do not present dichotomic (often isotropic) values due to present/ausent material, but a wide range of values due to the specific topology of the printed microstructure.

Topologically optimized designs ensure the optimization of a structural geometry regarding to certain requirements. The objective function depends on the variables that describe the problem. An extremal value of this function is pursued. There are designs to optimize the mechanical loads [4], the heat transfer [5] or to optimize the shape of the structure for the flux of a fluid around it [6]. The objective function usually only has one physical phenomenon to consider, but in the recent years some works couple more than one phenomenon. It is the case of Kruijf et al. [7] that studied in 2D generative designs to maximize the stiffness of the structure and minimize the resistance of heat dissipation. Guest et al. [8] used the topological optimization to design the microstructure of a small cube of an idealized material, to maximize the stiffness and fluid permeability. To fulfil manufacturing requirements, Lin et al. [9] used the Solid Isotropic Material with Penalization (SIMP) methodology to couple 2D generative designs and the multi-objective optimization method in order to make more realistic designs. And more recently, Querin et al. [10] were capable of using a multi-material approach in 2D generative designs through homogenization theory.

In optimizing the elastic energy density in the volume, there are mainly three ways of addressing the optimum design problem, so the optimal geometry does not have regions with low solicitations [11]. The differences are given by the approach used by each algorithm to reach the optimum design; namely to eliminate the parts of the structure with a low solicitation, to insert holes in the structure to optimize the geometry and to define the surface of the generative design. Although the final results can be very similar in many cases, the methodologies Evolutionary Structural Optimization (ESO) and Bi-directional Evolutionary Structural Optimization (BESO) [12] pose a heuristic criterion based on the natural evolution of the structure toward its optimal, by deleting the elements of a Finite Element mesh representing the mechanical behavior of the structure that show a lower level of stresses during successive iterations. BESO allows for the reintroduction of elements in the areas of the mesh with high stresses to relax them. Both methodologies are simple to implement and give good results with a low computational cost, but due to their structure, those seem not to be optimal and general enough to account for functionally graded anisotropic nonlinear metamaterials.

Another methodology based on the distribution of the material like ESO and BESO is the SIMP method [13]. In this method, the dependency between the isotropic properties of the material and the design variables are expressed in terms of the density of the material using a quadratic interpolation law. This makes the methodology less sharp than ESO and BESO where the elements are directly removed, but at the same time adds complexity to the calculation. Finally, the most complex methodology for the generative design of structures is the Level-Set method [14], which is arguably more satisfying from a mathematical point of view. In this method, a topological entity that represents the surface of the structure is defined, and a sensitivity analysis of the variation of this surface and its gradient to reach the geometrical optimum is performed.

In these methods, the “density” acts conceptually as an intermediate artificial variable, for example to determine the “amount” of material or to establish a limit in the optimization process (for example that only half of the volume contains material). But noteworthy, the density is not a relevant mechanical property in the common static analyses (if weight is neglected as an action), and indeed the assumed relation between the density and the mechanical properties does not have to hold in general. If we consider metamaterials, the density has very weak relation to the actual mechanical properties of the resulting continuum material. Indeed, many efforts have been directed to create methodologies to be added to the topological optimization to relate this artificial density with the real mechanical properties of the material, by using machine learning and coupled models at different scales [15, 16]. A deeper comparison between optimization methodologies can be found in Yago et al. [17]. Those works propose a new approach to consider spatial stiffness variability to define both the geometry of the generative design at the macro level, and the homogeneous stiffness of the material at every point to be fulfilled by the microstructure. This is done through the optimization of the elastic energy of the structure with a direct local variation of the mechanical properties.

## 2. The formulation of the topology optimization problem

Topology optimization based on density has two approaches whether the density is treated as a continuum field variable or a discrete field variable. In the first case, the optimization problem is written as

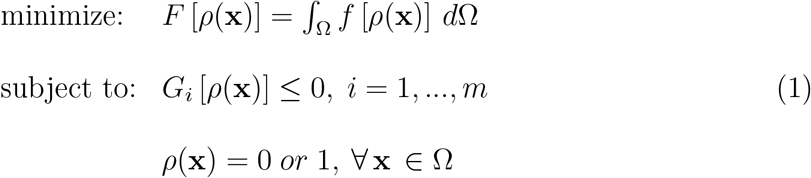

where *F* [*ρ*(**x**)] is the objective function subject to *m* different constraints *G*_*i*_, and the density *ρ*(**x**) is the continuum field variable (i.e. any location **x** may have its own value). These densities can take either value 0 (void) or 1 (solid material) at any point in the design domain Ω. Note that manufacturing procedures and design criteria can introduce additional constraints to the general problem. Besides, a volume constraint has been introduced in the literature with an academic purpose, as a basic constraint which improves general convergence.

In the second case, ***ρ*** denotes the design variable vector of length *N*, and *ρ*_*i*_ = *ρ*(Ω_*i*_) are its components, where *N* is the number of finite elements. This formulation establishes that the design variables *ρ*_*i*_ can only take discrete values: 0 or 1 within each element. Then, the optimization problem is written naturally in terms of the discrete problem as

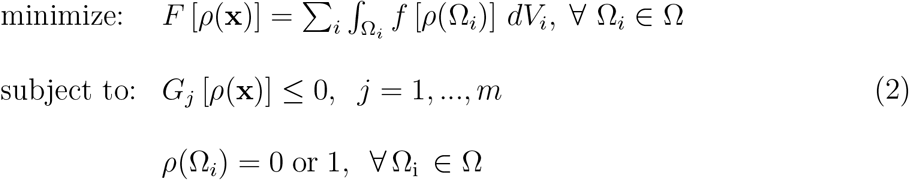

Whereas the discrete nature of the design variable is a typical difficulty in the mathematical treatment of the problem, there are several approaches capable of solving simple discrete problems with high efficiency.

On the other hand, because of different performance and manufacturing constraints non-dichotomic density design variables are often required. In such cases, the continuous topology optimization can be stated as in (2) but with a nonbinary value of *ρ*_*i*_, i.e. 0 ≤ *ρ*_*i*_ ≤ 1. Then, considering already a small strain linear-elastic equilibrium constraint, the optimization this optimization problem is

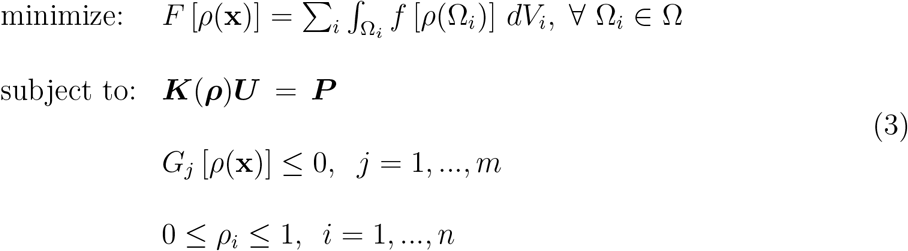

where *ρ*_*i*_ is the vector of density design variables, ***U*** is the displacements vector, ***K*** is the global stiffness matrix, ***P*** is the load vector and *G*_*j*_ represent the extra problem constraints. This approach constitutes the basis of the most successful optimization approaches: density methods [18], topological derivative [19], level set [20] and phase field [21].

One of the fucntions *F* [*ρ*(**x**)] that is typically minimized is the elastic energy. Under prescribed forces, the minimization of the elastic energy means increasing the stiffness of the structure. The elastic energy is

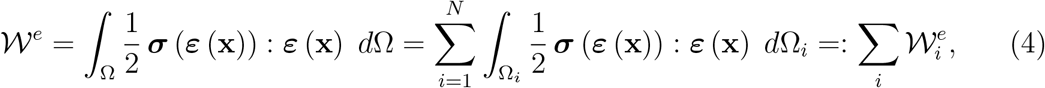

Of course, in the linear case, the elastic energy may be computed directly from the stiffness matrix and the displacements vector as

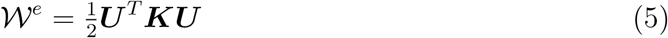

Shortly after the introduction of the first topology optimization methods, Bendsoe [18] and later Zhou and Rozvany [22, 23] proposed the so-called SIMP method, which is probably the most used optimization approach in industry. It is also similar to other used methodologies such as Rational Approximation of Material Properties (RAMP). In these methods, it is common the use of a penalty constant *π*. This constant is used in the update of the Young modulus, and also needs an optimization over the density variable. The difference between the SIMP and RAMP techniques is this update, namely

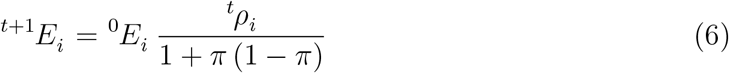

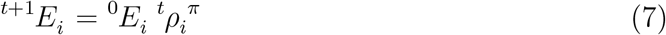

Examples are given in [24] for the SIMP approach, and in [25] for the RAMP approach.

As it is seen in the previous formulations, the density plays a major role in the procedures as an intermediate variable (note that we are dealing with static problems where weight is not considered). Indeed, the relations in Eq. (6) and Eq. (7) between “density” and Young moduli do not hold in general (except in the limiting cases), so this density should be considered just as an artificial variable.

In the following section we introduce a novel approach that does not use the density as intermediate artificial variable, but deals directly with the mechanical properties. In this work we deal with isotropic linear elastic materials, but since we deal directly with energies and meaningful material constants, the idea is powerful to be extended for nonlinear materials and to anisotropy. Moreover, the proposed approach directly creates a structure with graded variations of stiffness so its application combined with functionally graded metamaterials is more direct than with other methodologies [26].

## 3. The updated properties model (UPM)

The proposed approach is designed to minimize the total elastic energy of the full volume with the imposition of Dirichlet boundary conditions to restrict the displacements at some given locations, and Neumann boundary conditions to load the sample. Hence, our optimization problem is written as

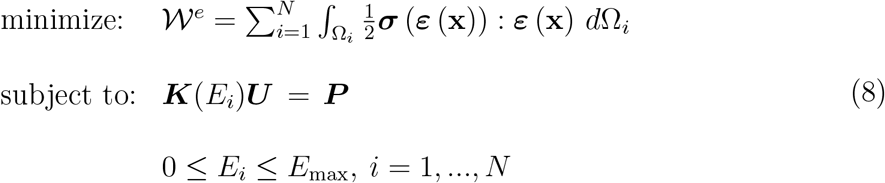

where *E*_*i*_ ≡ ***E***(Ω_*i*_) is the Young modulus of element *i*, the objective function is the structural elastic energy, 𝒲^*e*^. *E*_*i*_ are the design variables and *E*_max_ is the maximum Young modulus value (for example corresponding to the base printed material), necessary to achieve the convergence of the method. Hence, we operate directly with the Young modulus of each of the *N* elements of the volume.

Now we minimize the elastic energy iteratively without a gradient based approach, and without relying on the artificial parameter *ρ*. Since we are dealing now with linear elastic materials, we can write the stress tensor in terms of the Lame parameters as

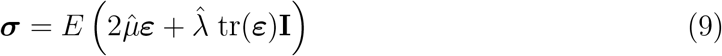

where *λ* and *μ* are the Lame parameters and 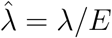 and 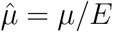 are the dimensionless counterparts. In the finite element context, the elastic energy is

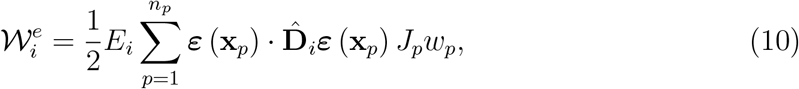

where *n*_*p*_ are the number of integration points considered, 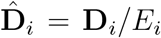 (being **D**_*i*_ and *E*_*i*_ the constitutive stiffness tensor/matrix and the Young modulus of an element *i*, respectively), *J*_*p*_ is the Jacobian of the element respect to normalized coordinates at the integration point *p*, and *w*_*p*_ are the weights for the quadrature. Since *E*_*i*_ is considered homogeneous in the element, 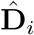 is constant—having the Poisson ratio *ν*_*i*_ fixed, as well. Therefore, the scalar *E*_*i*_ may be factored out.

The energy of an element *i*—given by Equation (10)—is now divided by the volume of an element, obtaining an average specific energy of an element 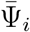, defined as

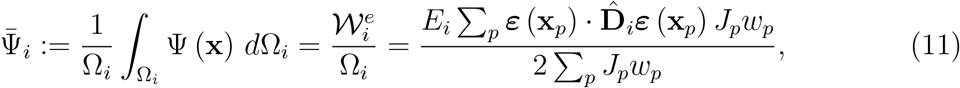

where 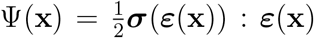. Note that the volume of an element is Ω_*i*_ = Σ_*p*_*J*_*p*_*w*_*p*_. Proceeding that way, the algorithm avoids the sensitivity regarding the different sizes of the elements of the mesh. Thus, the update is performed based on the importance of the specific energy of an element rather than the total elastic energy of the element.

We define the following strain variable:

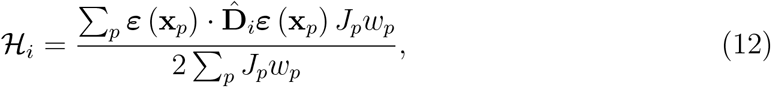

so the elastic energy of each element of the FEA discretization is:

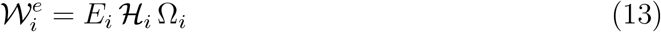

For elements with a unique integration point, i.e. *n*_*p*_ = 1, the previous expression simplifies into

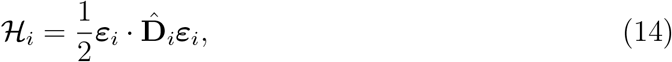

where ***ε***_*i*_ represents the strains tensor/vector (if in Voigt notation) evaluated at the unique integration point of the element *i*. We propose the following update properties model (UPM) to minimize the problem defined in (8):

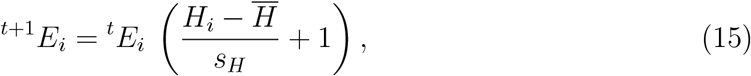

where 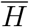 and *s*_*H*_ are the mean value and the standard deviation, respectively, considering all the *N* elements of the domain Ω. This update properties algorithm continues for every step *t* until convergence. Convergence is obtained when at two consecutive evaluations of the global elastic energy 𝒲^*e*^, we have

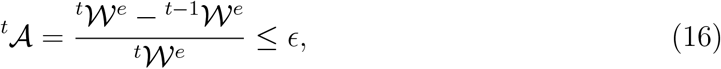

where *ϵ* is chosen tolerance close to zero.

The computational procedure of the previous equations is implemented in Algorithm 1. With this algorithm, the optimization step of the SIMP method is not needed, i.e. the approaches compare as:

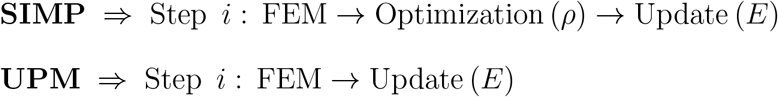

In the following examples the method has been implemented with a finite element formulation using 4-node thetrahedral elements and one integration point.

### Algorithm 1 Elastic energy minimization

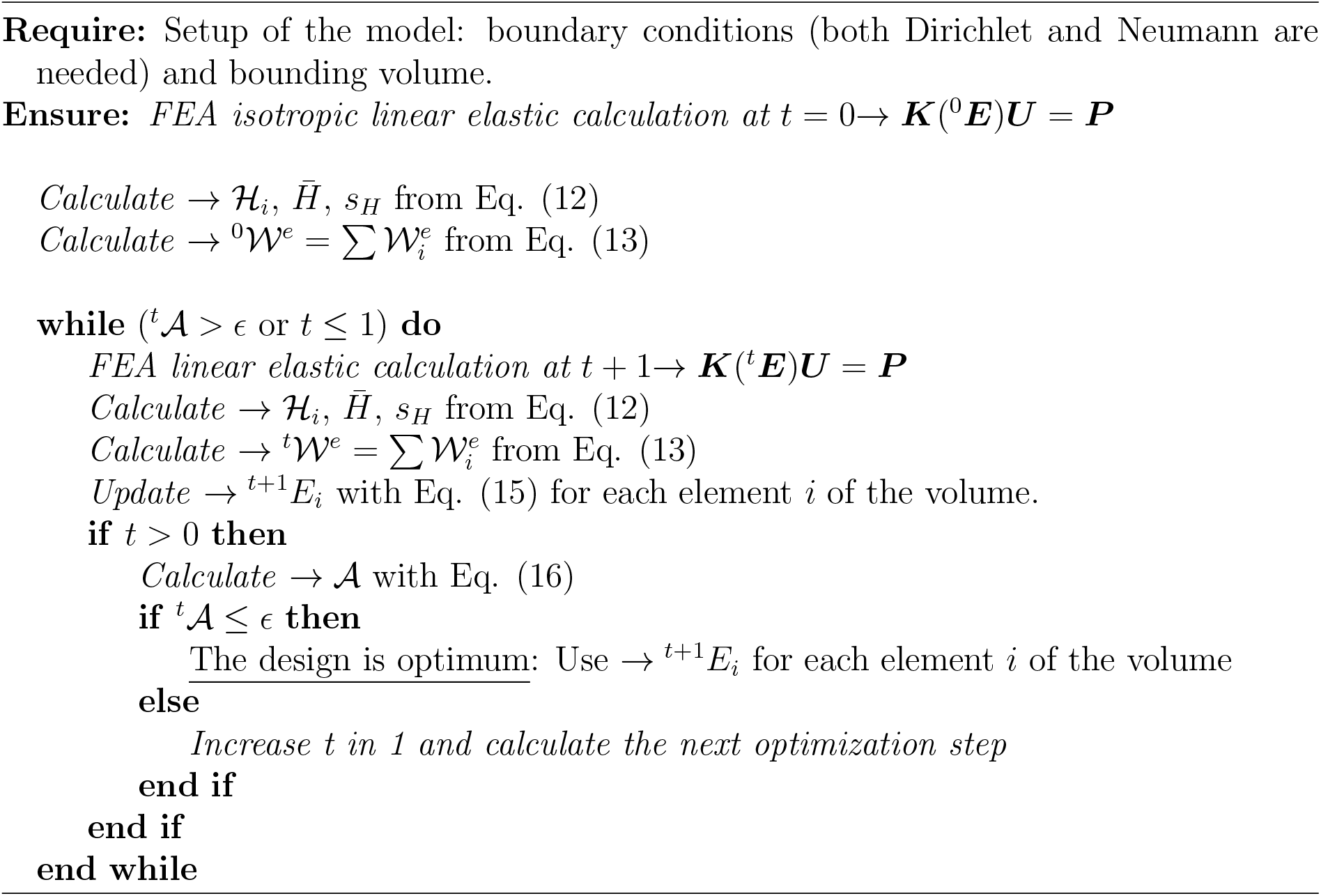

## 4. Examples

In order to test the algorithm presented, three different demonstrative, intuitive load cases using the same mesh have been analysed. The algorithm is designed for problems with a Dirichlet boundary condition for the restricted movements, and Neumann boundary conditions for the applied forces.

### 4.1. Loading cases

The loading cases are designed to cover different scenarios in terms of overall stress type and path. A first load case is intended to produce a stress field mainly influenced by normal tension. A second load case tests the capabilities of the model under a high shear state. A third load case focuses on a more general load and optimization case through the simultaneous use of different and non connected boundary conditions.

The calculations in all three cases are performed on a cube of 100 × 100 × 100 mm^3^ with 40000 linear tetrahedral elements, 9261 nodes and an element size of 5 mm. The displacements are restricted in all directions in the nodes of the blue circles of the Figure 1, and a load of 10 kN is imposed in the red circles in the upper side of the cubes, as shown in Figure 1.

**Figure 1:**
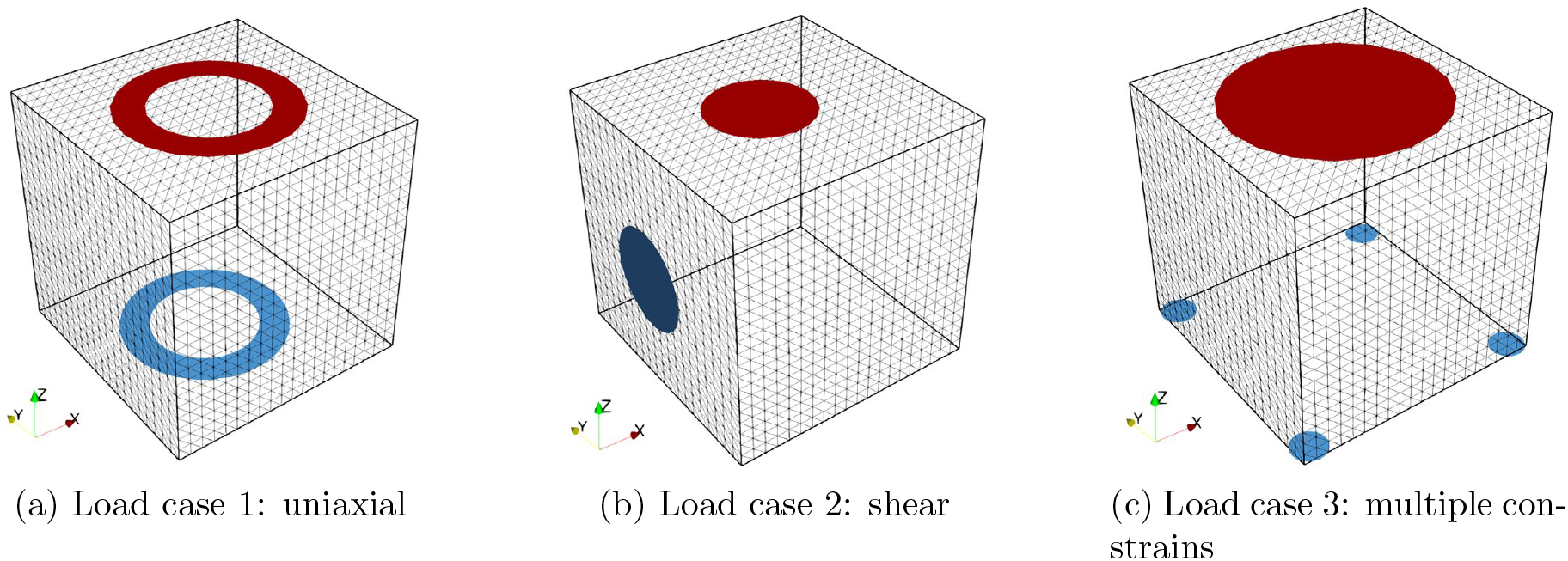
Mesh used and boundary conditions of the 3 cases studied. In the blue regions the displacements are restricted through Dirichlet boundary conditions. And in the red regions a vertical positive force is applied in each node through Neumann boundary conditions.

For each load case the particular boundary conditions are imposed as follows:

Load case 1: All the displacements are restricted in the blue disk of Figure 1a, located at *z* = 0. This disk has an outer radius of 35 mm and a inner radius of 25 mm. The same disk, but located at *z* = 100 mm and in red, is used to impose the forces. In each node a force of 10 kN is applied in the *z* direction.

Load case 2: All the displacements are restricted in the blue disk of Figure 1b located in the face of the cube at *x* = 0 mm. The disk is solid with an external radius of 20 mm. The same disk in red is used in the face of the cube at *z* = 100 mm to apply a force, along the *z* axis, of 10 kN in each selected note.

Load case 3: This case has multiple boundary conditions, the displacements are restricted in 4 blue solid disks of 5 mm radius, located each one in a different corner of the face of the cube at *z* = 0 mm, as Figure 1c shows. The force of 10 kN is applied to each node within a circle of 44 mm radius located at *z* = 100 mm, in red in the figure.

Figure 2 shows the strains field induced in the cubic sample when elastically loaded with an uniform Young modulus of 200 GPa. Figure 2a.1 and 2b.1 are the load case 1 under strain *zz* and shear strain *xz* (according to the axis of Figure 1). In this calculation the strain *zz* is concentrated around both the Dirichlet and Neumann boundary conditions, with an homogeneous distribution in the central region of the sample. The shear strain of this load case is insignificant in all the sample. Figure 2a.2 and 2b.2 corresponds to the load case 2. Here the shear strain is highly significant around the Dirichlet boundary condition, and the strain *zz* is more significant around the Neumann boundary condition, with even compression in certain regions of the sample. Figure 2a.3 and 2b.3 corresponds to the load case 3. This loading produces a strain *zz* concentration around both the Dirichlet and Neumann boundary conditions, with a higher concentration where the displacements are restricted. Again, the shear deformation is insignificant in this calculation, except for the node at the corner because its displacement is not restricted.

**Figure 2:**
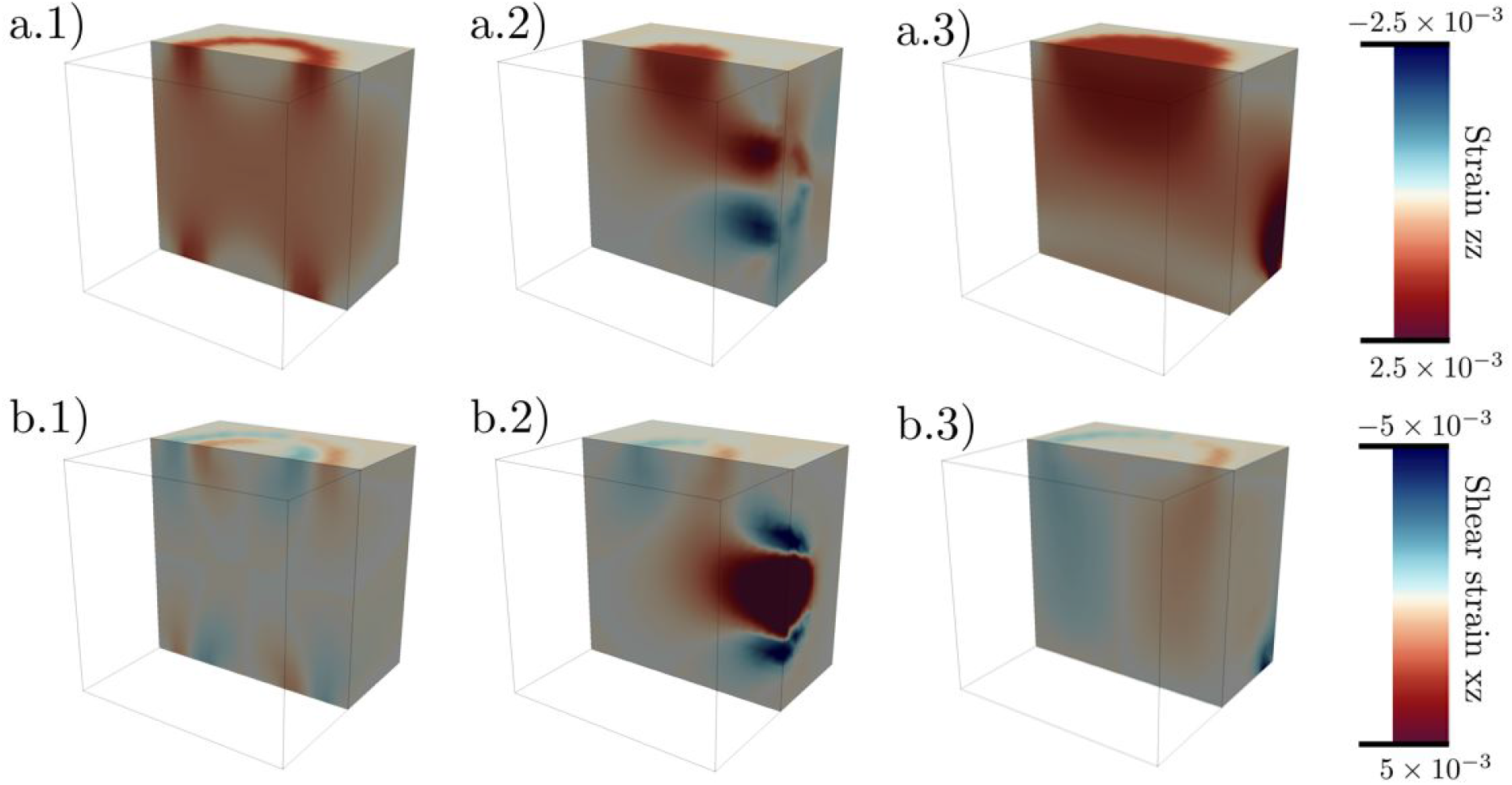
Vertical and shear strain fields for the 3 loading cases. Load case 1 (a), load case 2 (b) and load case 3 (c).

With those configurations, the convergence of the elastic energy at a minimum value is achieved within a small number of iterations of Algorithm 1, as Figure 3 shows.

**Figure 3:**
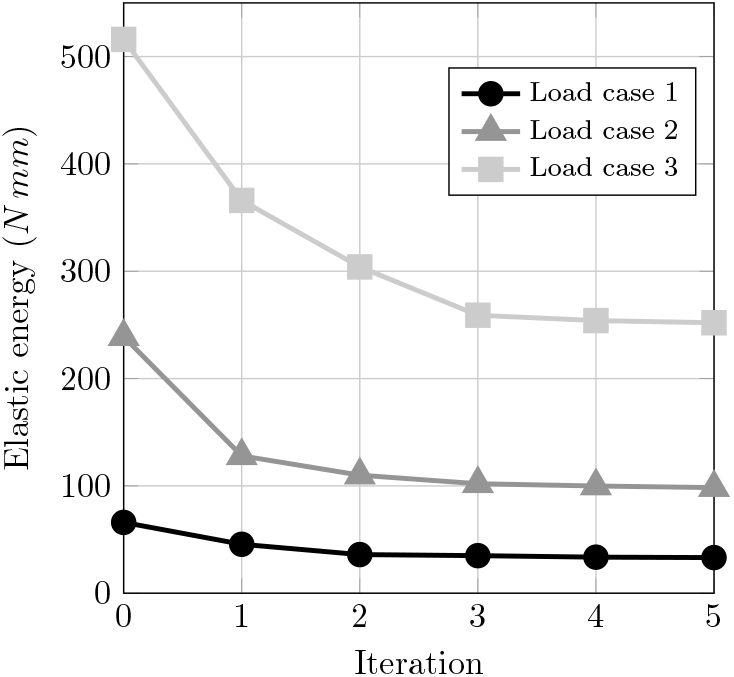
Elastic energy evolutions for the different iterations of Algorithm 1 for the three loading cases.

Figure 3 shows how the highest decay of the elastic energy is achieved within the first iteration of the algorithm, and the convergence is reached smoothly for the different loading conditions. In the next sections each loading case will be studied more in detail.

#### 4.1.1. Load case 1: uniaxial

The setup of this load case is shown in Fig. 1a. This has the same geometry for the restrictions of the displacements at the bottom and for the imposition of the forces on the top. The boundary conditions are imposed in two rings with the purpose of reproducing a uniaxial tensile load in a tube, so imposing the displacement restrictions and the loads applied, a tubular shape should be generated with the algorithm.

At the beginning of the calculation (Iteration 0), the cube is homogeneous and all the elements have the same Young modulus, assigned as 200 GPa. As Figure 4a shows, the values of the Young modulus are redistributed to reach the optimal design and the convergence of the elastic energy. So, as the iterations progress there are more elements with a low Young modulus (i.e. without structural influence and that could be eliminated if a lower threshold is established), and the elements with higher stiffness are increased.

**Figure 4:**
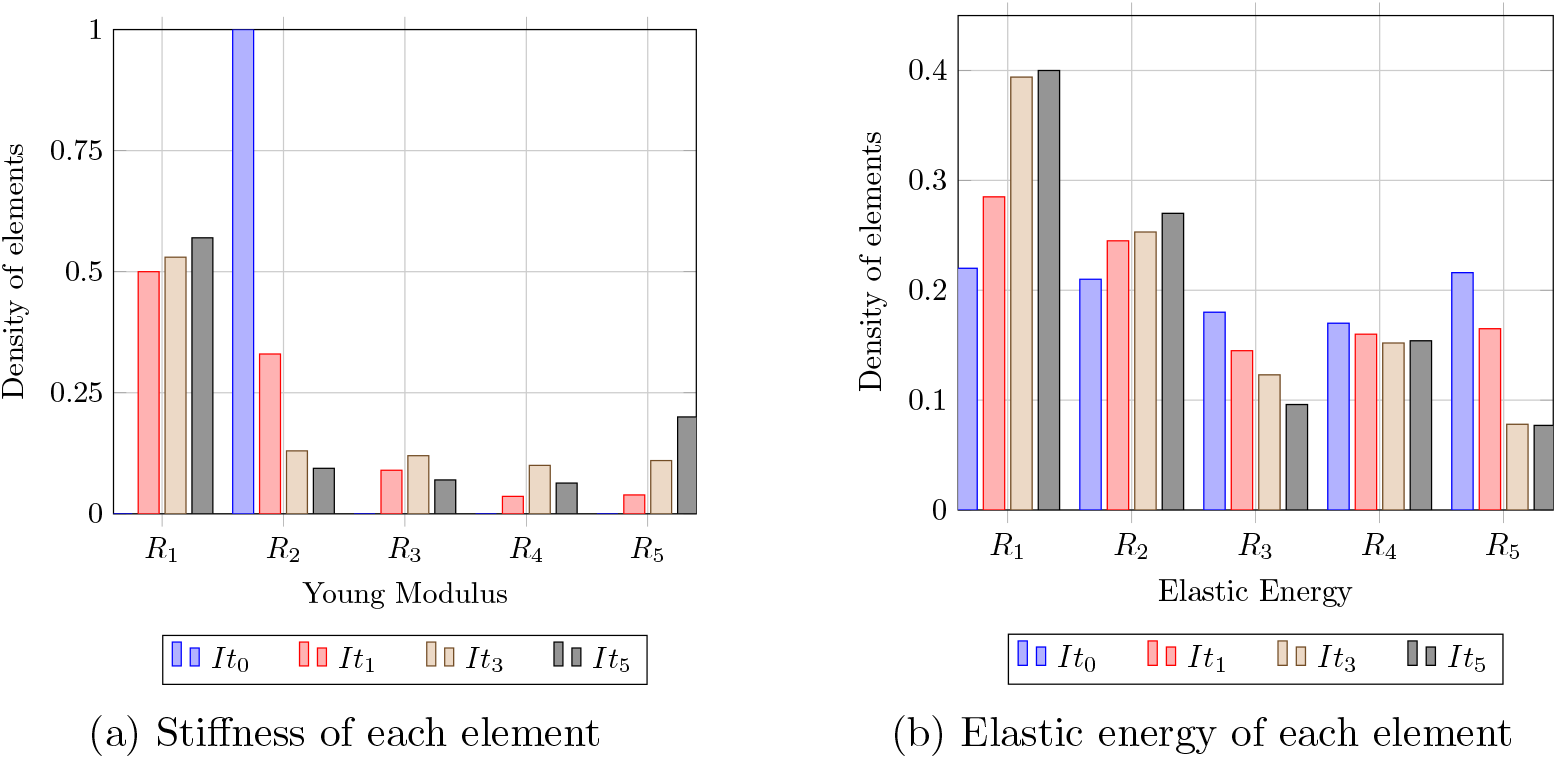
Load case 1. Redistribution of properties in the elements. In *a*) *It*_*i*_ represent the different iterations, and *R*_*i*_ the ranges, where *R*_1_ = [0, 150) GPa, *R*_2_ = [150, 250) GPa, *R*_3_ = [250, 350) GPa, *R*_4_ = [350, 450) GPa and *R*_5_ = [450, 500] GPa. In *b*) *It*_*i*_ represent the different iterations, and *R*_*i*_ the ranges, where *R*_1_ = [0, 1) Nmm, *R*_2_ = [1, 4) Nmm, *R*_3_ = [4, 7) Nmm, *R*_4_ = [7, 10) Nmm and *R*_5_ = [10, ∞) Nmm.

Figure 4b shows the variation on the distribution of the elastic energy stored by each element at each iteration. In the first iteration the elastic energy is roughly equally distributed due to the loading conditions in the mesh with an homogeneous Young modulus. As the iterations progress, there are more elements with a low elastic energy (i.e. without structural implications) and the proportion of elements with a higher elastic energy decrease.

Figure 5 shows the evolution of the structural generation through the different iterations and from different representations. The first column shows how the original cube is deformed with the boundary conditions, pulling from the top ring. And, as the stiff core is generated, the overall deformation decreases in the sample.

**Figure 5:**
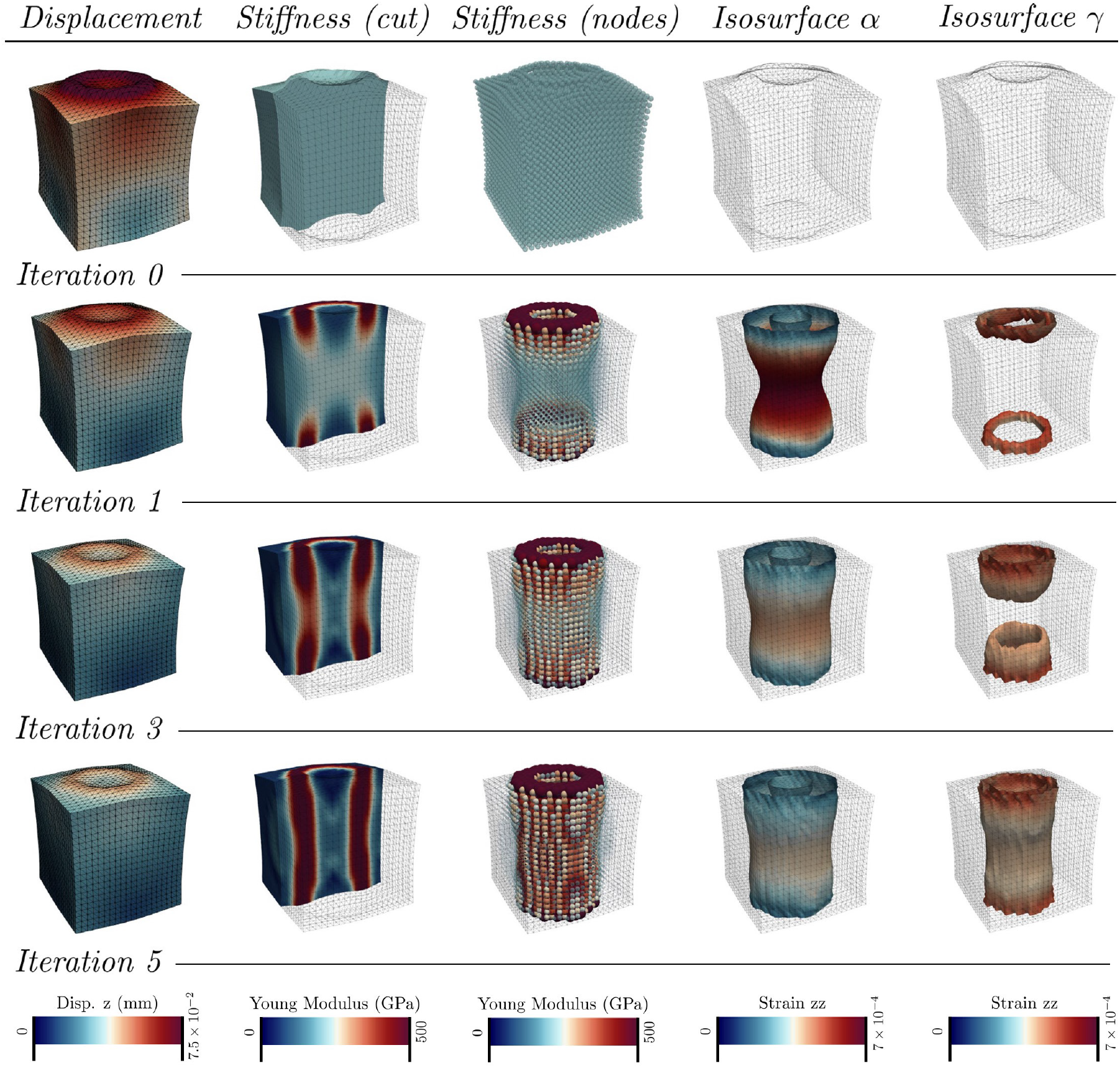
Evolution of the load case 1 along the different iterations. In the view of the stiffness (nodes), the size of the spheres is scaled with the value of the Young modulus at the node. The isosurface *α* is created by the elements with *E* = 200 GPa, and the isosurface *γ* is created by the elements with *E* = 450 GPa.

At iteration 1 the structure starts to be generated from the boundary conditions, with the growth of the stiff core, until both ends join at iteration 3, and the final geometry of the tube is defined at iteration 5, when the elastic energy converges totally. The strain *zz* is plotted in the columns with the evolution of the isosurfaces *α* and *γ*, and it shows how the strains are diminished during the generation of the structure.

#### 4.1.2. Load case 2: shear

This load case is configured by the restriction of displacements at a lateral circle, and the load imposed in another circle at the top. It creates a shear field.

Figure 6a has the same characteristics presented for Figure 4a, where the Young modulus is initially uniform during the first iteration and the initial configuration. But after Iteration 3 the Young modulus is redistributed, with a significant number of elements with a low stiffness. The number of elements with a high stiffness increases until the convergence of the calculation. Figure 6b presents the variation of the elastic energy, with an increment of the elements with a low energy (i.e. without structural capabilities), and a reduction of the number of elements with a high strain energy. Here the elastic energy of the elements is more concentrated outside the middle range, because of the loading conditions.

**Figure 6:**
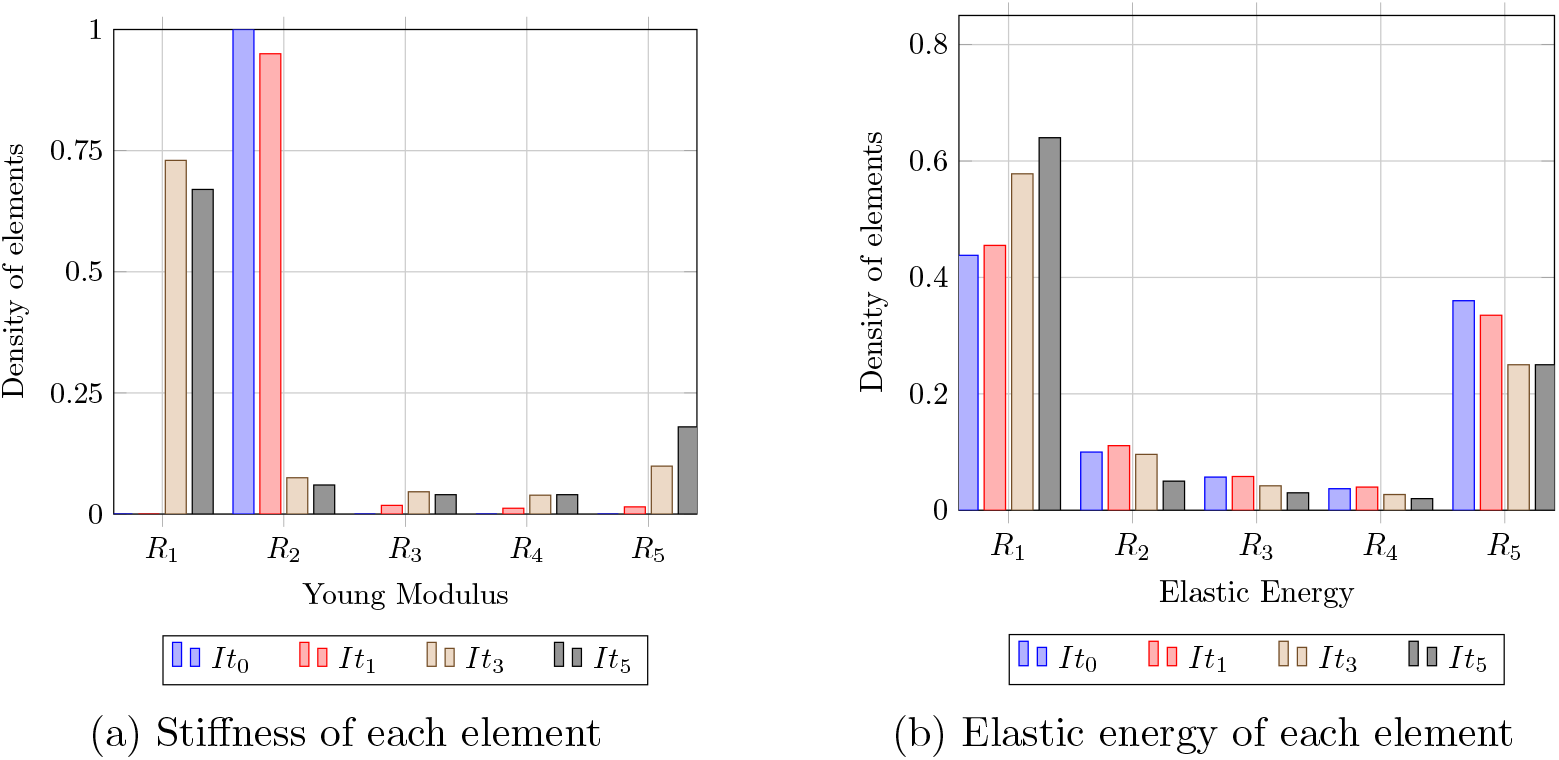
Load case 2. Redistribution of properties in the elements. In *a*) *It*_*i*_ represent the different iterations, and *R*_*i*_ the ranges, where *R*_1_ = [0, 150) GPa, *R*_2_ = [150, 250) GPa, *R*_3_ = [250, 350) GPa, *R*_4_ = [350, 450) GPa and *R*_5_ = [450, 500] GPa. In *b*) *It*_*i*_ represent the different iterations, and *R*_*i*_ the ranges, where *R*_1_ = [0, 0.5) Nmm, *R*_2_ = [0.5, 1) Nmm, *R*_3_ = [1, 1.5) Nmm, *R*_4_ = [1.5, 2) Nmm and *R*_5_ = [2, ∞) Nmm.

Figure 7 shows the behaviour of the sample during the generative design. The first column represents the deformation of the specimen, and the rotation induced by the loading conditions. As the stiff core is generated, this deformation decreases, with a reduction as well of the shear strain. The stiff core starts growing from the nodes with the Dirichlet boundary conditions to connect with the incipient core generated around the nodes with the Neumann boundary condition. Finally, this core gets thicker to reach the final configuration, when the equilibrium of the elastic energy is reached.

**Figure 7:**
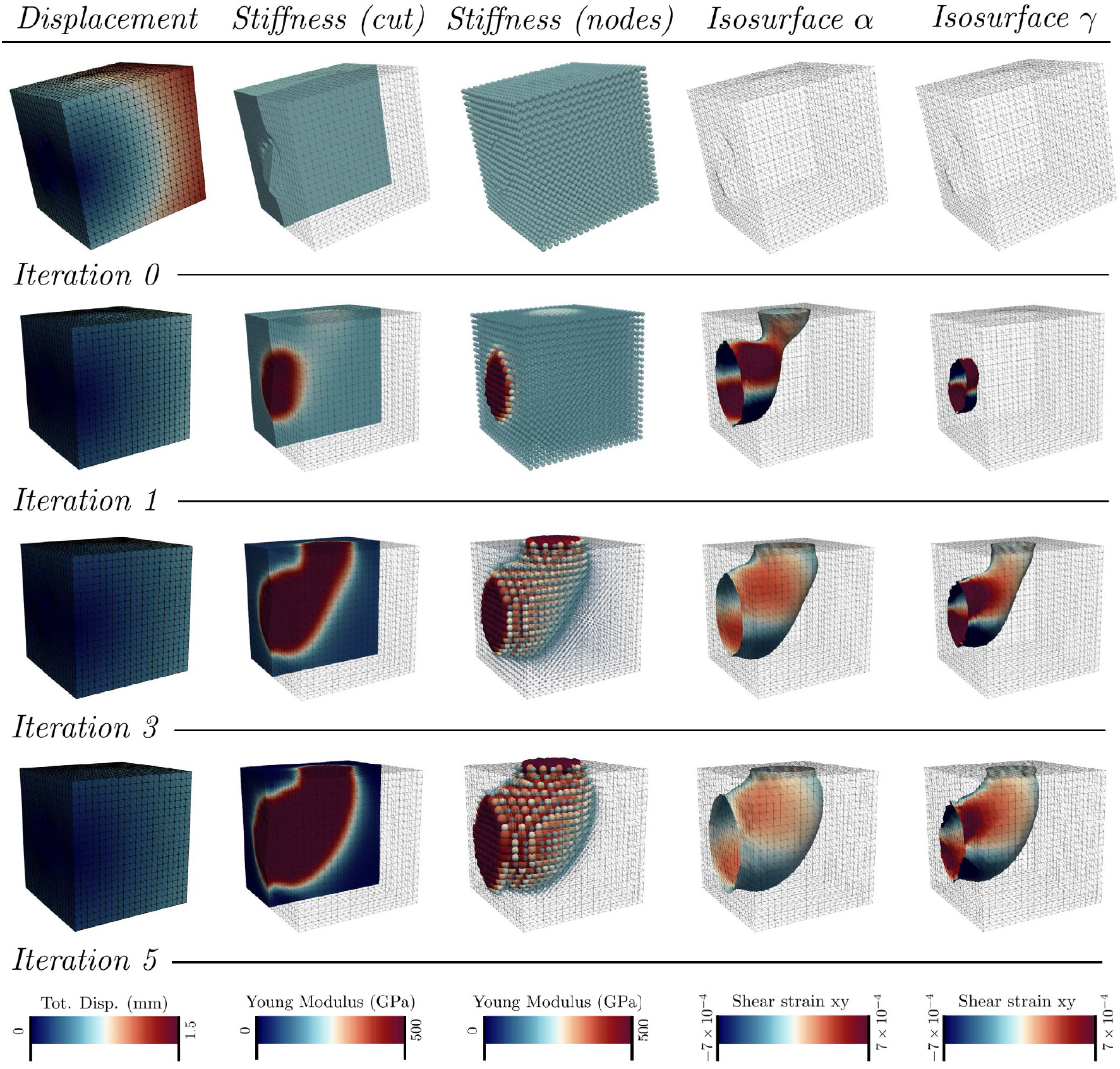
Evolution of the load case 2 along the different iterations. In the view of the stiffness (nodes), the size of the spheres is scaled with the value of the Young modulus at the node. The isosurface *α* is created by the elements with *E* = 200 GPa, and the isosurface *γ* is created by the elements with *E* = 450 GPa.

The shear strain is plotted in the isosurfaces, showing a decrease of this strain during the generation in the isosurface *α*. In the isosurface *γ* the high shear strain is maintained, and those elements are the ones with a higher elastic energy.

Finally, in Figure 7, the optimized design is an ‘elbow’ that connects the Dirichlet and Neumann boundary conditions. It can be represented with the isosurface at 200 GPa.

#### 4.1.3. Loading with multiple constrains

For load case 3, Figure 8 represents the evolution of the stiffness and elastic energy of all the elements through the different iterations. Here both the Young moduli and the elastic energy are shared along the ranges used. This configuration is more complex and the range of the elemental elastic energies is wider.

**Figure 8:**
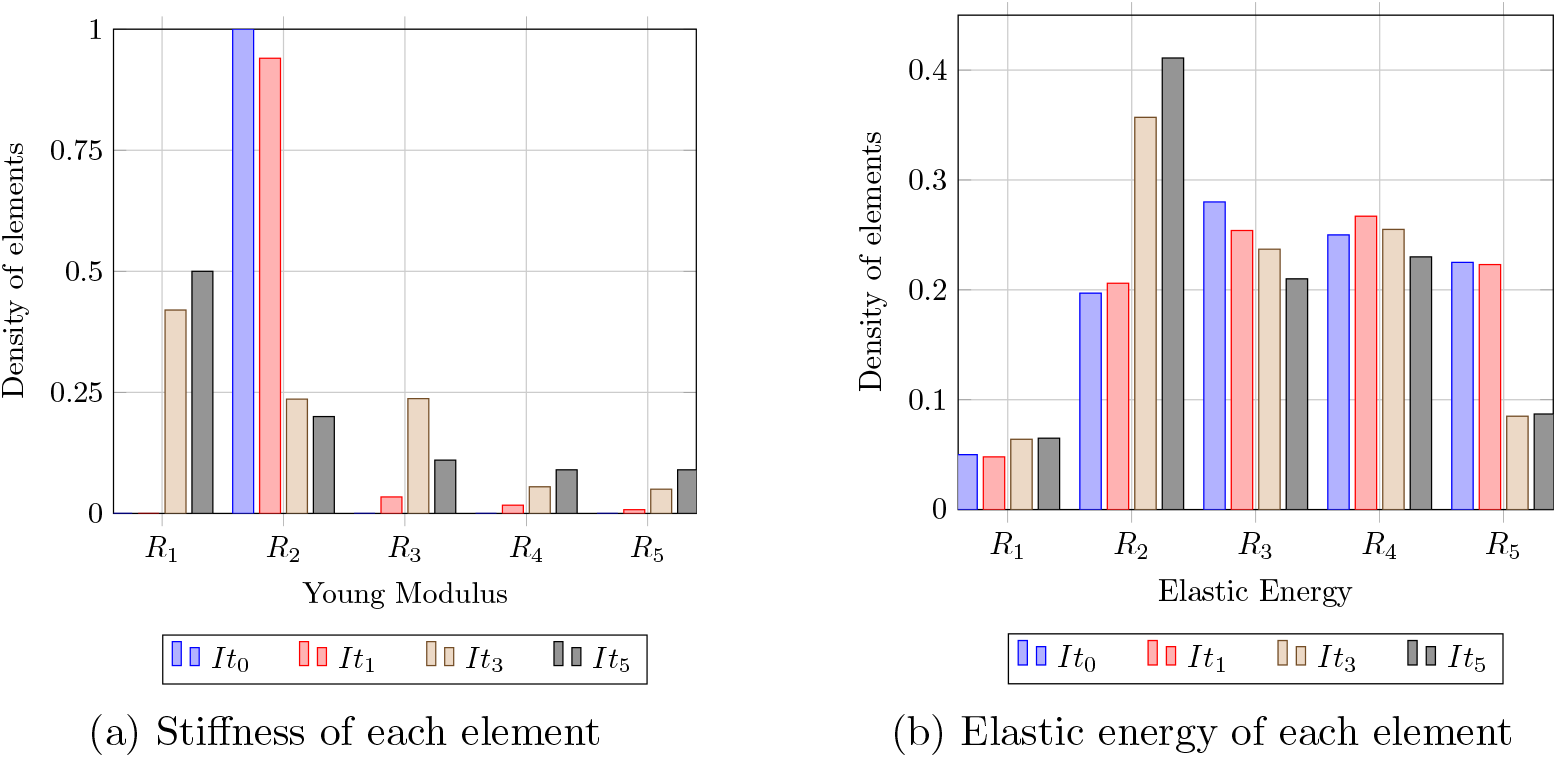
Load case 3. Redistribution of properties in the elements. In *a*) *It*_*i*_ represent the different iterations, and *R*_*i*_ the ranges, where *R*_1_ = [0, 150) GPa, *R*_2_ = [150, 250) GPa, *R*_3_ = [250, 350) GPa, *R*_4_ = [350, 450) GPa and *R*_5_ = [450, 500] GPa. In *b*) *It*_*i*_ represent the different iterations, and *R*_*i*_ the ranges, where *R*_1_ = [0, 1) Nmm, *R*_2_ = [1, 10) Nmm, *R*_3_ = [10, 25) Nmm, *R*_4_ = [25, 50) Nmm and *R*_5_ = [50, ∞) Nmm.

The evolution of the Young modulus within the nodal set of the sample is shown in Figure 9. As in the previous loading cases, the stiff core starts to be generated from the nodes with Dirichlet boundary condition, growing to connect with the stiff core created around the nodes with the Neumann boundary conditions. Finally, the optimized design creates a structure with 4 legs (typical of a stool) that minimizes the elastic energy of the structure.

**Figure 9:**
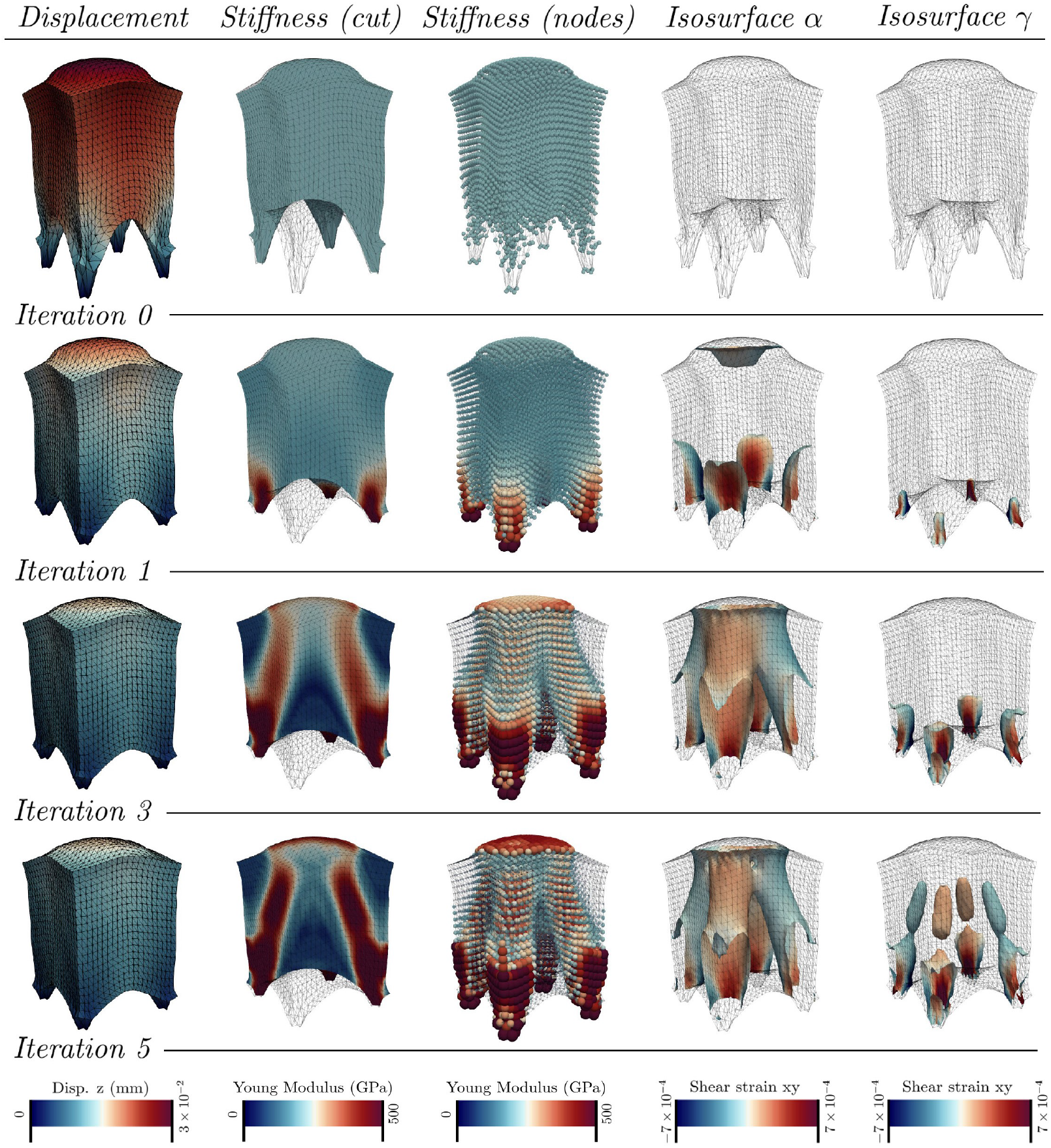
Evolution of the load case 3 along the different iterations. In the view of the stiffness (nodes), the size of the spheres is scaled with the value of the Young modulus at the node. The isosurface *α* is created by the elements with *E* = 200 GPa, and the isosurface *γ* is created by the elements with *E* = 450 GPa.

### 4.2. Comparison with the SIMP model

The optimization results from two representative load cases using our approach are compared with the optimized results from the SIMP methodology in Figures 10 and 11. Recall the different design variables employed in each method: the elementary Young modulus (*E*_*i*_) of the proposed methodology, versus the fictional densities (*ρ*_*i*_) employed in the SIMP method. The results from the SIMP method are obtained with the commercial software OptiStruct [27], with a penalty factor for the SIMP model of *p* = 2. We depict the density isosurfaces for this method. In our case, we use our own finite element code and depict surfaces of isomoduli. Whereas our computational times for these simulations are in the order of those used by OptiStruct (with differences smaller than 10%), we have not optimized the code (e.g. we did not use sparse solvers). Hence, we expect that with equal optimization, the code will be much faster, giving that we do not need to perform the optimization phase in the density.

**Figure 10:**
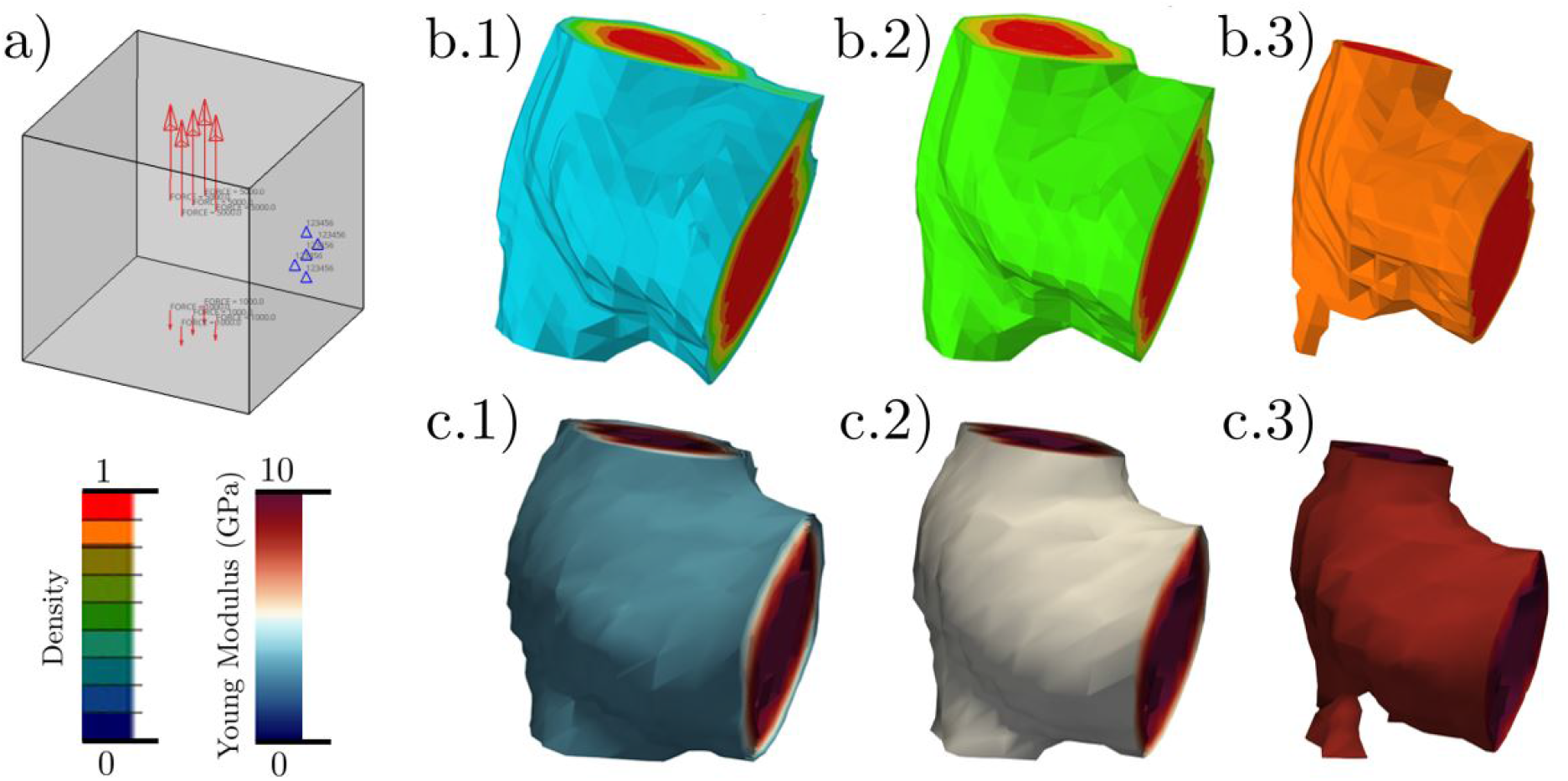
Comparison of the results from OptiStruct and the proposed method. Load case 1: a) Shows the loading conditions used and the color codes for the bandplots. b.1), b.2) and b.3) show the variation of the isosurfaces for different values of the density when using the SIMP method. c.1), c.2) and c.3) show isosurfaces of Young moduli using our method for Young moduli approximately equivalent to those of the SIMP method. The purpose of the comparison is to show that the shapes of the isosurfaces are similar in both methods.

**Figure 11:**
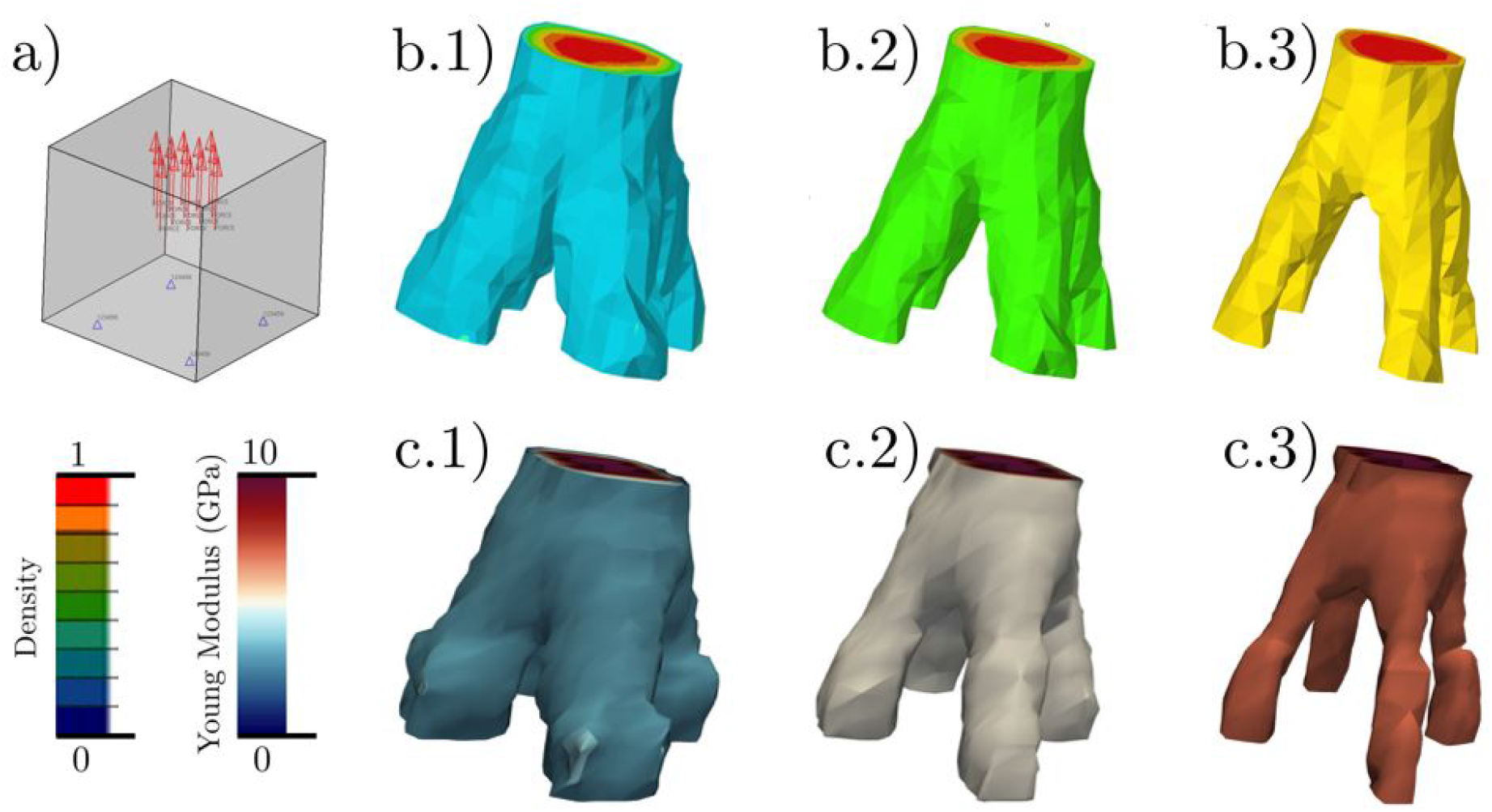
Comparison of the results from OptiStruct and the proposed method. Load case 2: a) Shows the loading conditions used and the color codes for the bandplots. b.1), b.2) and b.3) show the variation of the isosurfaces for different values of the density when using the SIMP method. c.1), c.2) and c.3) show isosurfaces of Young moduli using our method for Young moduli approximately equivalent to those of the SIMP method. The purpose of the comparison is to show that the shapes of the isosurfaces are similar in both methods.

In Figures 10 and 11 the following analogy can be used to compare both results: *E* = *E*_max_ is equivalent to *ρ* = 1, the same as *E* = 0 is equivalent to *ρ* = 0. For the values between these boundaries, the relationship between *ρ* and *E* is not trivial, but with some *ρ* the same contour generated by certain *E* values can be reproduced. Those Figures present the optimization result for each load case, where the stiffness and density distribution has been filtered following the previous analogy. As an approximation, if *E*_max_ = 10^5^ MPa and OptiStruct’s density result is filtered for *ρ >* 0.8, then, the stiffness layout shows values for *E >* 0.8 × 10^5^ MPa.

The Figures 10 and 11 show the final distribution and geometry achieved during the optimization. Note that these simulations are slightly different from the previous ones considered. In both figures 10 and 11, the *b* plots correspond to the SIMP results, with the density values given in the legends in Figs. 10a and 11b. The *c* plots in both figures correspond to the Young moduli isosurfaces obtained with our method. The selected isosurfaces are those which roughly correspond to the density values of the SIMP method, so the shapes of the isosurfaces from both methods can be visually compared.

However, although the results match our intuitive expectations, it is worth comparing them with those obtained by using commercial software, such as OptiStruct.

## 5. Conclusions

In this work we present a novel approach for topology optimization. Our approach consists on a direct update of the mechanical properties of the material, instead of the use of the density with a penalty factor optimized in an intermediate layer. Hence, we work directly with strain energies and mechanical properties, which in the linear case results equivalent to work with Young moduli. We have demonstrated that the optimized designs are intuitive and in line with the results obtained with the SIMP method

Working directly with the mechanical properties is not only arguably more efficient, but it also has advantages for concurrent structure-material design using functionally graded metamaterials, since for example, the metameterial may be designed to have continuous spectrum of elastic moduli which is not directly dependent on a density, but on a specific design of the microstructure. Functionally graded metamaterials is becoming an important field thanks to the use of 3D printing techniques, where the structure may be optimized both at the macro level (external geometry of the design) and at the micro level (variation of mechanical properties). Furthermore, working directly with energies and mechanical properties may result in extensions of the procedure to nonlinear behavior and anisotropy, aspects also inherent to designs using metamaterials. We are currently undergoing these extensions.

## Author Contributions

LSM: Conceptualization, methodology, writing original draft. IBY and HGM: Coding and examples. MAS: Laboratory management, supervision and editing. FJM: Funding acquisition, supervision, editing. All authors contributed to manuscript revision, read, and approved the submitted version.

## Funding

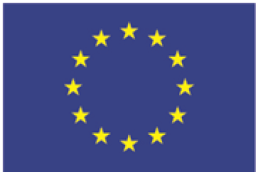

*This project has received funding from the European Union’s Horizon 2020 research and innovation programme under the Marie Slodowska-Curie Grant Agreement No. 101007815*

